# Managing Marek’s disease in the egg industry

**DOI:** 10.1101/431304

**Authors:** Carly Rozins, Troy Day, Scott Greenhalgh

## Abstract

The industrialization of farming has had an enormous impact. To most, this impact is viewed solely in the context of productivity, but the denser living conditions and shorter rearing periods of industrial livestock farms provide pathogens with an ideal opportunity to spread and evolve. For example, the industrialization of poultry farms drove the Marek’s disease virus (MDV) to evolve from causing a mild paralytic syndrome to causing a highly contagious, globally prevalent, disease that can have up to a 100% mortality rate. Fortunately, the economic catastrophe that would occur from MDV evolution has been prevented through widespread use of live imperfect vaccines that limit disease symptoms, but fail to prevent transmission. Unfortunately, the continued rollout of such imperfect vaccines is steering the evolution of MDV towards an even greater virulence and an ability to evade vaccine protection. Thus, there is a need to investigate alternative economically viable control measures for their ability to inhibit MDV spread and evolution. In what follows we examine the economic viability of standard husbandry practices for their ability to inhibit the spread of both virulent MDV and very virulent MDV throughout an industrialized egg farm. To do this, we parameterized a dynamic MDV transmission model and calculate the loss in egg production due to disease. We find that the MDV strain as well as the cohort duration had the greatest influence on disease burden and hence egg production. Additionally, we find that the standard husbandry practice involving conventional cages, often referred to as “battery cages”, results in the least per capita loss in egg production due to MDV infection when compared to alternative enriched or aviary (free-run) systems for virulent MDV, but not very virulent MDV, in which case the Aviary system performs the best. These results highlight an important cost that managers will face when implementing new hen husbandry practices.

## 1. Introduction

The industrialization of farming empowers farmers to keep pace with the ever-increasing demands of consumers, but it also creates a situation highly conducive to pathogen evolution as a result of cramped living conditions and shorter rearing periods (ANTHONY, 1998). A primary example of this is Marek’s disease virus (MDV), which causes Marek’s disease (MD), a disease of poultry that has evolved from a relatively harmless paralytic syndrome into a highly virulent pathogen (Witter, 1997) as a result of industrialization (Atkins et al., 2013a; Rozins and Day, n.d.). To make matters worse, MDV is highly contagious, globally prevalent (Dunn and Gimeno, 2013), causes up to 100% mortality (Read et al., 2015; Witter, 1997), and imposes a colossal economic burden (Morrow and Fehler, 2004).

The economic burden of MD comes from both direct losses from hen mortality and morbidity (e.g., egg production loss), and indirect losses caused by industry wide use of vaccines and control measures. While indirect losses are substantial, the control of MDV starting in 1969 through a succession of live vaccines has saved billions of dollars and helped to ensure the present economic stability of the industry (Churchill, AE and Payne, LN and Chubb, 1969; Churchill et al., 1969). Unfortunately, current vaccines for MDV have major drawbacks in that they only limit disease symptoms and permit both infection and transmission. Such drawbacks enable virulent MDV strains to go undetected and are attributed with being a major factor in the continued evolution of MDV virulence (Read et al., 2015), including its potential to evade vaccine induced immunity (Nair, 2005).

Detection and eradication of MDV is extraordinarily difficult. MDV spreads through freely circulating viral particles (Biggs P. M., 1967) that are shed through the feather follicles of infected laying-hens. As infected laying-hens are likely symptom free due to vaccination, and cohort sizes range from 30,000-100,000 hens (Holt et al., 2011), the identification and removal of MDV infected hens is difficult. Additionally it has been shown that MDV is often reintroduced to barns as often as once per month (Kennedy et al., 2018). Thus, other measures are needed to prevent or limit MDV infection.

Here, we evaluate the most common management scenarios in the egg industry for their ability to prevent MDV infection, while also mitigating any egg production loss. Specifically, we investigate how animal husbandry practices such as the density of laying-hens (reflecting alternative caging systems), the cohort duration, and the MDV strain influence MDV incidence, mortality, and egg production. Using data on MDV infection, in addition to data on the demographics of typical laying-hens in the egg industry, we evaluate the influence of management scenarios on MDV incidence, mortality, and egg production. We use a mathematical model (Rozins and Day, 2016) calibrated to reflect ongoing industrial practices, namely those of Aviary, Conventional, and Enriched systems. We assess the effect of MD on egg production over a 10-year horizon to evaluate any effect of management scenario and MDV infection on egg production.

## 2. Materials and Methods

To quantify the impact of management scenario on mitigating the effects of MDV infection on egg production, we use a mathematical model for MDV transmission in industrial poultry farms of laying-hens (Rozins and Day, 2016). As the vast majority of laying-hens are vaccinated (Payne, 1985), our model assumes full vaccination coverage with the current gold standard vaccine Rispen CVI988 (Ralapanawe et al., 2016a). We also assume two distinct mechanisms of MDV transmission: 1) transmission within a cohort of laying hens (those sharing a barn), and 2) transmission between consecutive cohorts of laying-hens occupying a barn (i.e. transmission through residual viral particles left in the barn).

### 2.1 Management Scenarios

We evaluate 72 different management scenarios. These scenarios describe the typical egg industry, matching flock sizes to the stocking densities of Aviary (30000 laying-hens), Conventional (80000 laying-hens), and Enriched (50000 laying-hens) systems, and 12 different cohort durations that capture the most common molting practices (no molt (NM), one molt (OM) at 69 weeks, and two molts (TM), one at 69 weeks and another at 104 weeks) (Bell, 2003). Molting is a natural process in which hens lose and regrow their feathers and briefly stop laying eggs. The process rejuvenates laying-hen production for additional cycles. We also consider a very virulent MDV strain (vvMDV) (pathotype FT158) and a virulent MDV strain (vMDV) (pathotype MPF57). For each scenario, we estimate the probability of a MDV epidemic, the flock mortality, and the average egg production over a 10-year horizon (Table S7, Fig.2, Fig. 3).

### 2.2 Egg production

To evaluate the impact of MD on egg production over a 10-year horizon, we base uninfected laying-hen egg production on a study of 25 million White Leghorns laying-hens (Figure 1) (Bell, 2003). MDV infected laying-hens incur a 5% egg production loss (*R*), which is an expected consequence of MDV infection (Purchase, 1985). Both uninfected and infected laying-hens’ egg production is continuously discounted by the annual US 2016 inflation rate of 1.25%. Further details of parameters, including sources, are available in Table 1.

**Figure 1.**
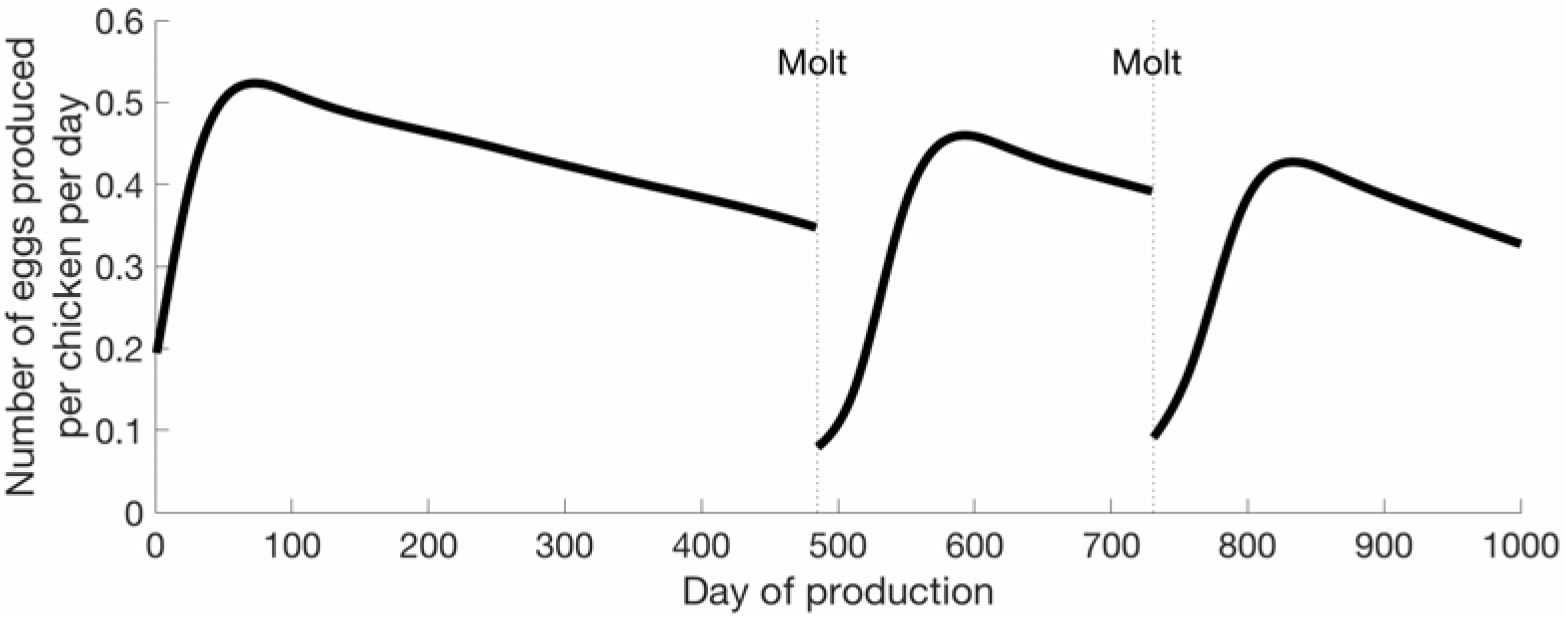
Mean daily egg production per hen over a 1000-day period. In this scenario, the hen is molted twice, once after 69 weeks (438 days) and again at 104 weeks (728 days). Molting rejuvenates the egg laying process, but has diminishing effects over time, and is done at most twice.

### 2.3 MDV transmission with-in a cohort

To describe MDV transmission within a cohort of laying-hens, we parameterize a compartmental model (Rozins and Day, 2016) to capture MDV prevalence levels in concurrent vaccinated flocks of laying-hens occupying a barn. We consider two classes of laying-hens: those susceptible to MDV infection (*S*), and those infected with MDV (*l*). The rate at which susceptible laying-hens become infected is governed by a constant transmission rate (*σ*), and the density of viral particles (*F*) in the barn. Infected laying-hens experience disease related mortality at rate (*ν*), and shed viral particles at the rate (*κ*). Finally, viral particles are removed, either through decay or the ventilation system, at rate (*δ*). Overall, the system of differential equations that models MDV transmission with-in a cohort is:

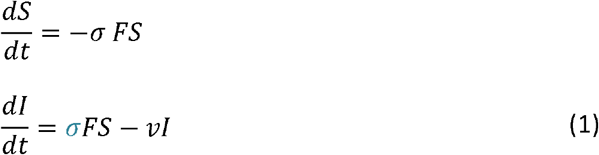

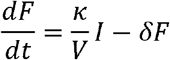

where *V* is the volume of the barn. Note, implicit in the definition of system (1) is the assumption that the per capita contact rate between susceptible laying-hens and viral particles is density dependant.

### 2.4 MDV transmission between cohorts

After a cohort duration of *T* days, the barn is emptied, cleaned, and restocked with the next cohort of new susceptible laying-hens. The emptying, cleaning, and restocking period is modelled as instantaneous, as the time to accomplish this is small relative to the duration of egg production. Thus, the laying-hens and density of viral particles *after* the farm is emptied, cleaned, and restocked is given by the difference equation:

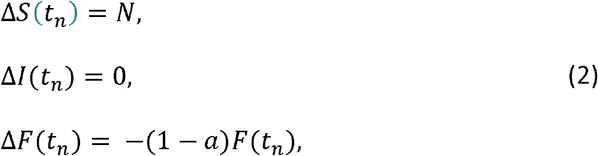

Where *N* is the flock size, *a* is the proportion of viral particles that remain after the barn is emptied, cleaned, and restocked, *t*_*n*_ = *nT* and *n* is an indexing for the cohort.

By combining the between cohort equations (2) and the within cohort equations (1), we have a model to describe the continual chain of transmission of MDV in cohorts of laying-hens over a 10-year horizon.

### 2.5 Parameter values

All model parameters were extracted from available data (Atkins et al., 2013a, 2011a; Cui et al., 2016; Kennedy et al., 2018; Ralapanawe et al., 2016b, 2016a; Zhang et al., 2015). Parameters associated to barn characteristics where obtained from communication with Burnbrae Farms in Ontario Canada, or the literature. We assume dust shed by MDV infected laying-hens contain viral particles proportional to the virulence level of the MDV strain (Ralapanawe et al., 2016a). To obtain the viral shedding rate, we took Atkins et al. (2011) model of daily dander shedding for a typical broiler bird and arameterized it according to a recent study on dust shed from MDV infected layer chickens vaccinated with Rispen CVI988 (Atkins et al., 2011a; Ralapanawe et al., 2016b, 2016a). Thereby, we obtain the viral shedding rates *κ* for each MDV strain from data on the dust shed from a typical laying-hen (Table S4), and the viral copy number (VCN) of MDV per milligram of dust for vMDV (pathotype MPF57) and vvMDV (pathotype FT158) (Table S3), and over each cohort duration (older chickens will shed more viral particles) (Table S5) (Bell, 2003; Witter et al., 1968). Data on the virulence level of the MDV pathotype and mortality, in addition to the standard assumption of Exponentially distributed parameters that is typical of compartmental models(Greenhalgh and Day, 2017; Hethcote and Tudor, 1980), was used to estimate MDV mortality rates (*ν*) for each MDV strain (Ralapanawe et al., 2016a). We base the viral particle removal rate (*δ*) on the barn ventilation system and estimates of the decay rate of the viral particles (Kennedy et al., 2018). Therefore, *δ* is taken as the sum of the decay rate and the average air exchange rate of a typical barn (Table 1, Webappendix). In practice, the air exchange rate of a barn is based on its stocking density, as more densely stocked barns require a greater exchange of air compared to less densely stocked barns. For simplicity, we consider a constant air exchange rate, which we estimate using available data and standard fitting techniques. Finally, we determine the transmission rate *σ*_*ν*_ using both an estimate, and the mathematical formulation of the MDV effective reproductive number (Atkins et al., 2013, 2012; Renz, 2008) (Webappendix), yielding

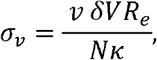

where *κ*, *δ*, *ν*, are previously estimated (Table 1, Table S4, Table S5).

### 2.6 Sensitivity analysis

To quantify the contribution of each model parameter to the variability of the outcomes measured, we calculated 95% confidence intervals and first-order sensitivity indices.

**Table 1.**
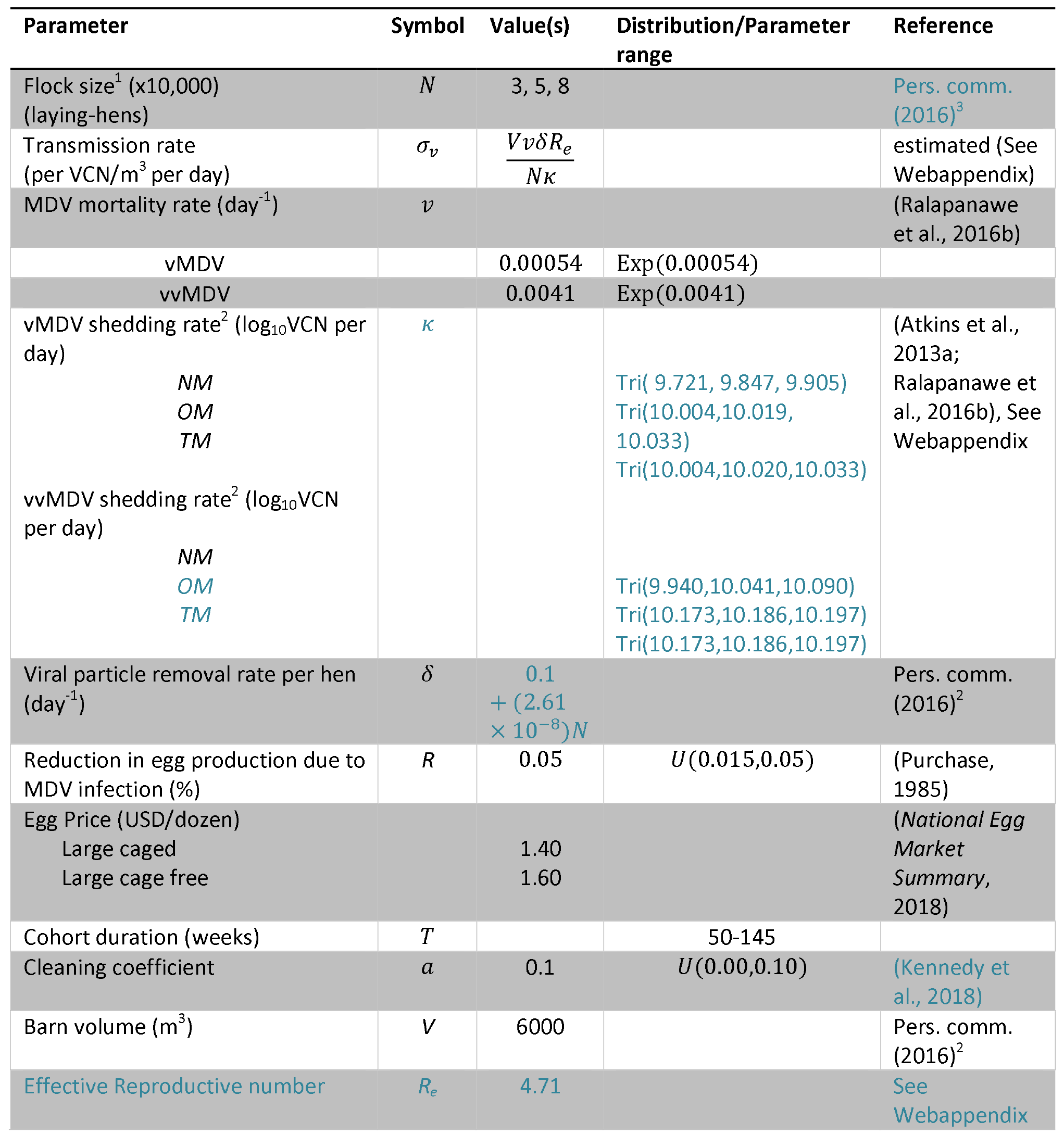
Model parameter values.

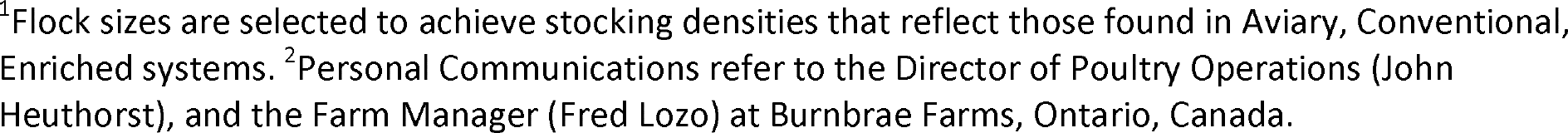

## 3. Results

We found that the strain of MDV circulating within the barn has the most significant impact on egg production. Additionally, the cohort duration (the time hens spend laying eggs) has an impact of egg production, with longer cohorts suffering greater losses. But perhaps most interestingly we found that barns with the greatest stocking densities suffered the least per capita loss in production when vMDV circulated within the barn, but when the more virulent vvMDV was circulating, the Aviary system (the least densely stocked barn) had the least per capita loss in egg production.

### 3.1 Disease

Our findings show that, once established, the majority of a flock will become infected, regardless of management scenario. We also found that disease spread was very rapid, infecting the majority of a flock well before the end of the cohort. Mortality due to MDV infection varied significantly across management scenarios and was largely dependent on the cohort duration and the MDV strain virulence level (Fig 2b, Fig 2c). The less virulent strain of MDV would often fail to become established within the barn (it was eliminated by the end of the first cohort), or would become eliminated before the end of the 10-year study period due to between-cohort cleaning (Fig 2a). The more virulent strain was more successful at becoming established within the barn (Fig 2a).

**Figure 2.**
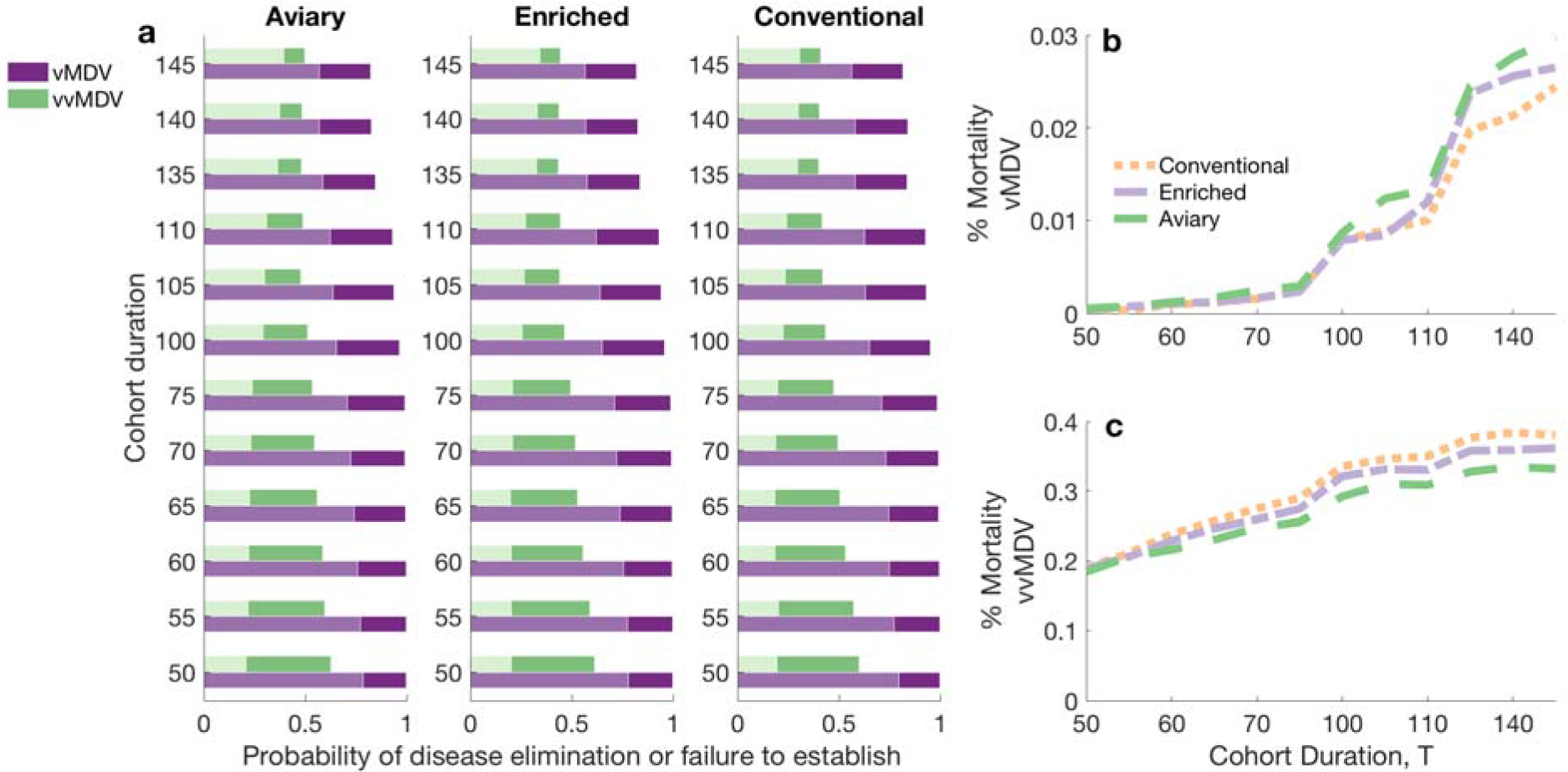
Likelihood of MDV elimination, or failure to become established after introduction, within a 10-year period, and MDV mortality. (a) Likelihood MDV is eliminated within 10 years for particular cohort duration, MDV strain, and system. In purple is the probability of disease elimination for a barn seeded with vMDV and in green for a barn seeded with vvMDV. The light shading (light purple or light green) indicates the proportion of time disease is eliminated by the end of the first cohort (after cleaning), otherwise the disease is eliminated over multiple cohorts. Mean mortality (mean proportion of the flock that die due to MDV) for each cohort duration due to (b) vMDV and (c) vvMDV for Aviary, Conventional, and Enriched system stocking densities.

### 3.2 Cohort duration and molting

Longer cohort durations experience higher mortality and greater per capita egg production loss due to MDV infection (Table S7 and Fig 2). This was greatest for the more virulent strain of MD (vvMDV). For example, laying-hens in a conventional system over a 65-week cohort duration (zero molts) had a mean mortality rate of 23% and produced on average 17.63 fewer eggs per hen when vvMDV was established. This egg production loss amounts to a $2 USD loss per bird, but in a conventional barn with 80,000 hens amounts to 10.15 million eggs and approximately $1.3 million (2018 USD) over 10-years. Under the same setup, if laying-hens are kept for an additional 40 weeks (undergoing one molt) the loss more than doubles to 39 eggs per hen, or $2.7 million over 10-years.

**Figure 1.**
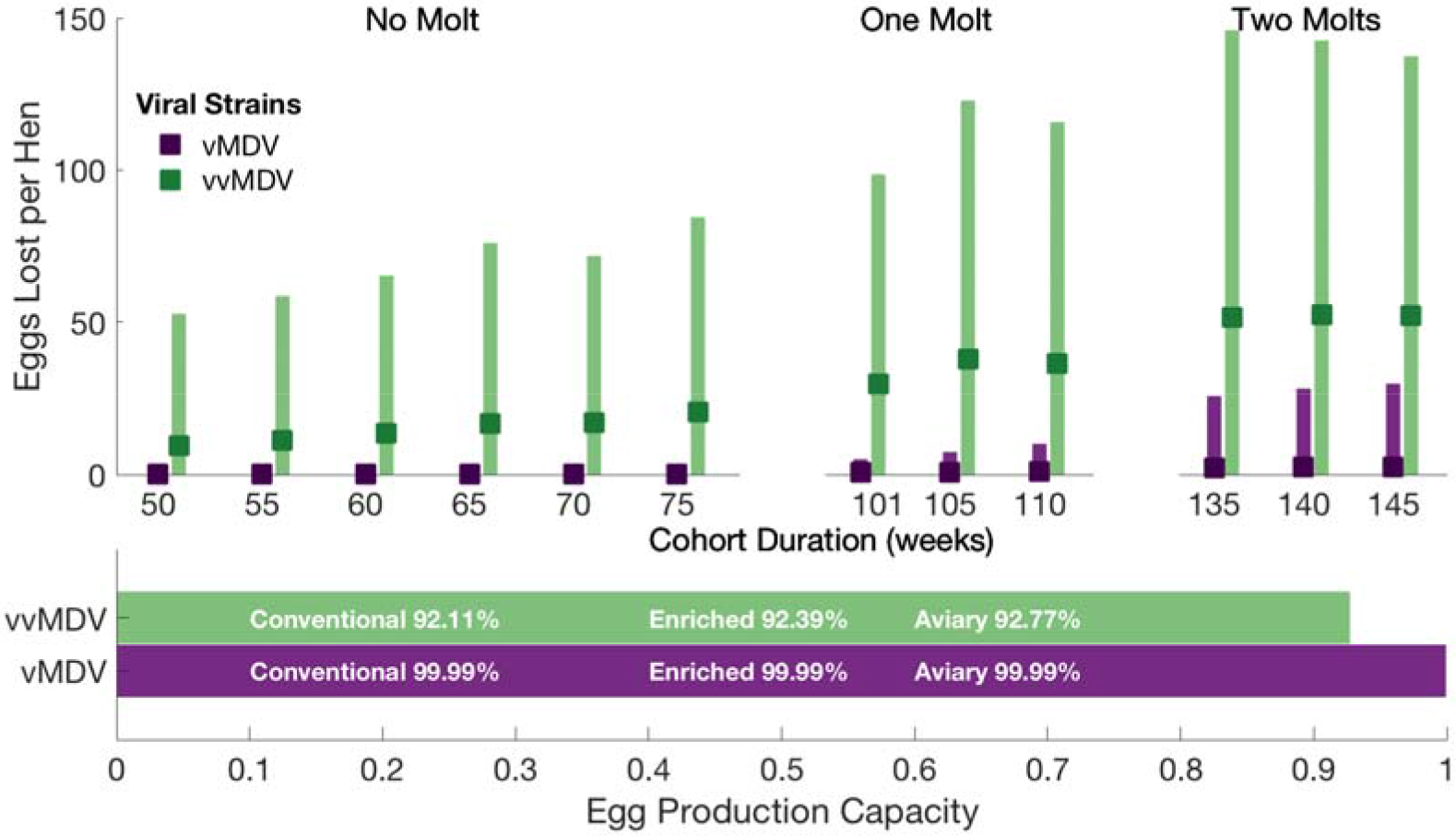
(Top) Average eggs lost per hen with 95% credible intervals for all cohort durations and molting practices (no molts, one molt, and two molts) and (Bottom) Egg production capacity for a 60-week cohort, measured as (eggs produced/total possible eggs producible in the absence of disease) for each MDV strain and Aviary, Conventional, and Enriched system stocking densities.

**Figure 2.**
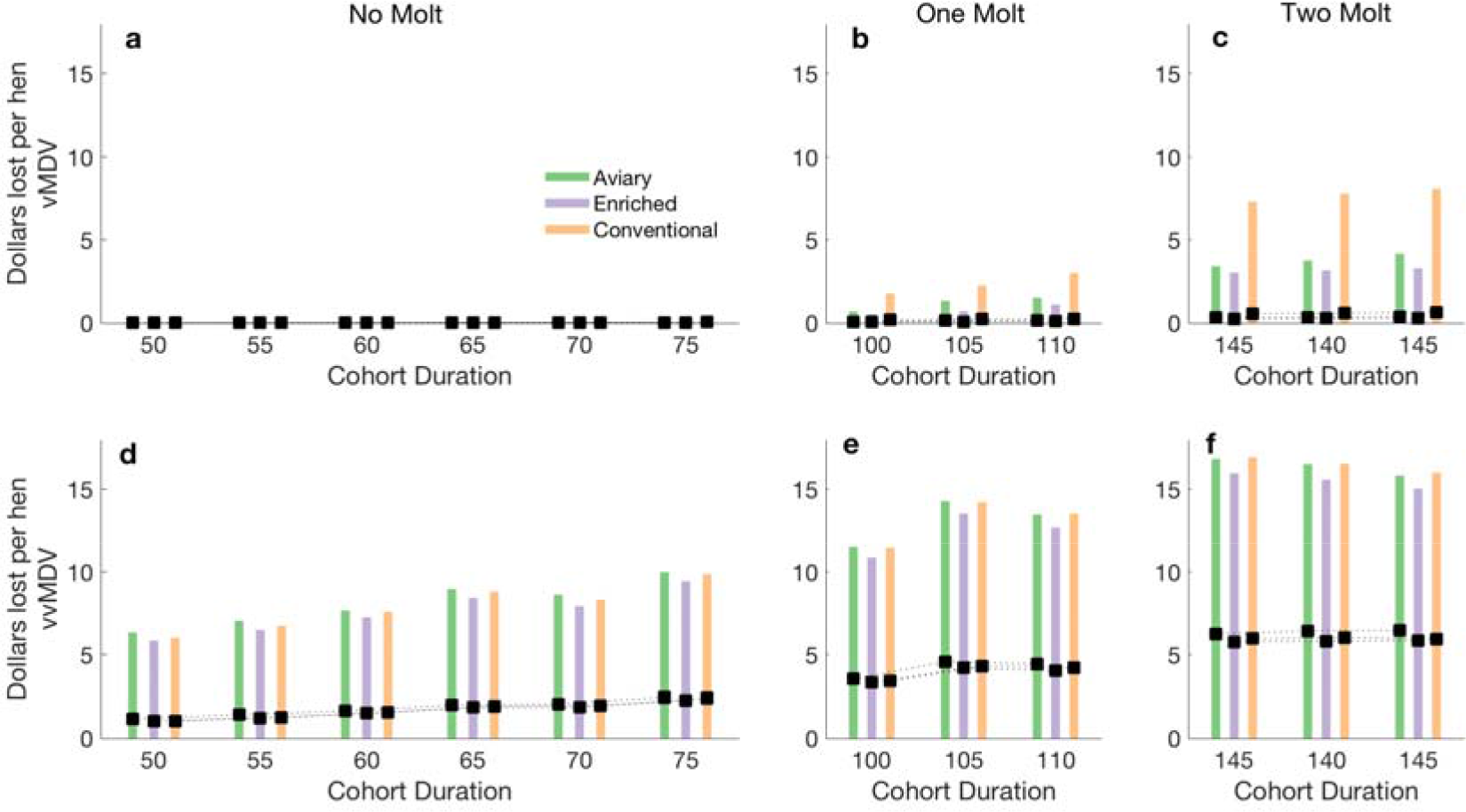
Economic loss (USD), due to mortality, and reduction in egg production due to MD. Results are scaled by stocking density and future egg-earnings are discounted using the 2016 US inflation rate of 1.25%. To calculate the total loss in USD, we first calculate the total revenue possible for a scenario (Table 1) and then subtract the total revenue when MDV is introduced. Finally, to normalize across stocking densities, we divide by the total number of laying-hens. Plots a-c) examine the vMDV strain, and plots d-f) examine the vvMDV strain. Plots a) and d) reflect a barn that does not molt its hens (no molting), plots b) and e) a single molting and plots c) and f) two moltings. The bars represent the 95% credible interval while the points represent the mean.

### 3.3 Stocking Density

Aviary, Conventional, and Enriched systems had similar per capita egg production losses. However, switching from a Conventional system to an Aviary or Enriched system yielded a small per capita egg production loss of less than one egg per hen when the less virulent strain on MD (vMDV) was circulating (Table S7). However, for the more deadly strain of the virus (vvMDV), switching from a Conventional system to an Aviary system yielded a mean per capita gain of approximately two eggs per hen on average and a switch to an Enriched system yielded a mean gain of approximately one egg per hen (Table S7). Note that the overall losses were greatest for the Conventional system (more hens more losses) when we accounted for hen numbers, the per capita loss in the conventional system was always less than that in the Aviary system for vvMDV but not vMDV.

### 3.4 MDV Strain

The circulating MDV strain had the greatest effect on laying-hen survival and egg production (Fig. 2b, Fig. 2c, Fig. 3, Table S7). The vvMDV strain caused at least twice the per capita egg production loss of the vMDV strain (Fig. 3). For both MDV strains, the per capita egg production loss increased as cohort duration increased (Fig. 2).

### 3.5 Sensitivity Analysis

To quantify the contribution of parameters to the variability in predicted egg production, we calculated variance-based first-order sensitivity indices (Sobol, 2001) (Table S6). First-order sensitivity indices indicate how uncertainty in a particular parameter contributes to the variability of model outcomes. Details of the probability distributions used in the calculation of first-order sensitivity indices are available in Table S6. Predictions in egg production were most sensitive to the MDV mortality rate *ν* and the reduction in egg production due to MDV infection *R* (Table S6). Sensitivity to MDV mortality increased with cohort size and cohort duration, while the opposite relation was observed for the reduction in egg production due to MDV infection (Table S6).

## 4. Discussion

This study is the first to evaluate the burden of MDV on the most common management scenarios in the egg industry using mathematical models. Additionally, the mathematical model developed is also the first calibrated to the Rispens CVI988 vaccine (Rispens et al., 1972), the current gold standard vaccine in the poultry industry. Given these firsts, we evaluated the effects of two MDV strains on laying-hen production in Aviary, Conventional, and Enriched systems, including the typical laying-hen cohort durations for a 6000 m^3^ barn. For each scenario, we estimated MDV prevalence, the potential for disease elimination, and the egg production loss (measured per capita) due to MDV infection.

While theory suggests less densely stocked barns should make disease elimination easier [13], we observed MDV infection persisting for all stocking density scenarios. In all scenarios, if the outbreak persisted, disease spread was rapid, with the majority of hens infected by the end of a cohort. The high prevalence is consistent with other modelling studies of MDV transmission within broiler farms that featured significantly shorter cohorts durations (Atkins et al., 2013b; Kennedy et al., 2018). The lower laying-hen numbers in Aviary systems carry an additional disadvantage, as lower stocking densities and more open designs render flocks more prone to hen cannibalism (Ahammed et al., 2014). It also leads to higher per capita egg production losses from MDV infection for the less virulent strain of the disease (vMDV) and a larger per capita financial loss in production for all but the shortest cohorts used in this study. However, the Aviary system had the least per capita loss in egg production when the more virulent strain (vMDV) was circulating within the barn. Additionally, lower laying-hen numbers should make barn cleaning easier, as there are fewer hens to shed viral particles, and thus fewer accumulated viral particles at the end of a cohort to clean, implying a greater likelihood of interrupting transmission of MD between cohorts.

Financially, the more densely stocked Conventional and Enriched systems performed the best. This is due to a number of factors. The caged eggs produced in the Conventional and Enriched systems are of less value than the free-run eggs of the Aviary system. Therefore, even when the Aviary system had a lower per capita loss in production, the overall per capita financial loss in the Aviary system was always greatest. Additionally, for shorter cohorts there is a diluting effect for the more densely stocked barns, as there are more hens to infect, which takes slightly longer in these barns. Therefore, by the end of the cohort a smaller proportion of hens, in the more densely stocked barn, have become infected or dies due to disease. Finally, the higher air exchange rate, which occurs at higher stocking density, removes infected MDV particles at faster rate. Combining this result with lower hen cannibalism rates, a 13%-36% lower upkeep cost (Matthews and Sumner, 2015), and the fact that higher stocking densities have been shown to select for less virulent strains of MDV (Rozins and Day, n.d.), show that Enriched and Conventional systems have substantial utility in the fight against MDV. However, since MD spreads rapidly throughout a barn, Conventional systems can have up to 80,000 hens shedding viral particles accumulating within the barn throughout the laying period. Therefore, if eradicating MDV from a barn is the objective (rather than managing symptoms through vaccination), then a barn using a Conventional system may require more cleaning than a barn stocking fewer hens.

Mathematical models inevitably involve simplifying assumptions. For instance, our model treated all eggs produced as equal value and quality. This assumption understates the productivity of longer cohort durations, as older hens produce larger eggs that are often worth more. Furthermore, we did not account for the increased cost associated to the 47% more hens required over a ten-year period for husbandry practices that avoid molts (Bell, 2003). We also did not account for multiple factors that may affect MDV persistence, such as transmission from direct contact of laying-hens in open environments, the effects of littler floors, which is a known factor affecting hen welfare and disease transmission (Dawkins et al., 2004; Lay Jr et al., 2011), seasonally-varying ventilation rates, or potential breaches in biosecurity. Incorporating such additional modes of transmission would likely strengthen the utility of Conventional systems, as multiple modes of transmission would make elimination more difficult. In addition, we do not account for the effects of disease coinfection, although with additional compartmentalization we could develop co-infection models with various other diseases. Additionally, all parameter values used in this model were constant with respect to time. Previous modelling efforts of MDV transmission in the broiler industry (Atkins et al., 2013a, 2013b; Kennedy et al., 2018) have assumed parameters such as transmission and the viral shedding change over time. We believe many of the parameter values, had we modelled them as functions of time, would asymptote quickly, and since the lifespan of a layer hen is much longer than that of a broiler bird, it is unlikely that assuming constant parameter value has a significant impact of the overall results of the model. Finally, while parameterizing our model we used information derived from studies on broiler birds (Atkins et al., 2011b) when information was not available for laying hens, and we are unsure of the consequences, if any, this has on our results. With this in mind, the methods used to construct our impulsive compartmental model could also easily be applied to study other avian diseases, such as the highly pathogenic H7 avian influenza recently found in a Tennessee commercial flock (Karlsons, Donna and Cole, 2017), as the impulse feature of the model naturally describes the all-in-all-out dynamics of poultry farms.

For the scenarios our model considered, our results illustrate that the Conventional and Enriched systems are best for mitigating per capita losses when the less virulent strain, vMDV, is circulating, but the Aviary system does best for mitigating per capita egg loss when the more virulent strain, vvMDV, is circulating within the barn. This is likely due to the higher stocking densities which requires additional time for the same proportion of the flock to become infected. This leads to higher production for the slower spreading vMDV strains. Additionally, the higher ventilation rates of the Conventional and Enriched systems help to delay the onset of the MD outbreak in a cohort. However, due to the additional cost of free-run eggs, the Aviary barn suffers the highest per capita financial loss in egg production for all scenarios and both viral strains explored.

## Conclusion

With the fear that prolonged use of Rispens CVI988 is masking the emergence of extremely virulent MDV strains (USAHA, 2015) and so enhancing current husbandry practices to combat MDV is of the utmost importance. Our results show that natural disease elimination through regular management practices is unrealistic for the more virulent vvMDV. Our results also suggests that increasing stocking densities helps to reduce per capita egg production loss due to MD in the short term (for the less virulent strains) and helps to mitigate long-term virulence evolution (Rozins and Day, n.d.). While at first these results may seem surprising, they call to light that improving long-term hen welfare is not as simple as solely reducing stocking densities or cohort durations, and that the improvement of hen welfare is not necessarily distinct from the goals of economics.

## Author’s Contributions

SG and CR came up with the study design and they parameterized the model. All authors participated in the drafting and editing of the paper.

